# A Model Quantitative Assessment Tool for Nonhuman Primate Environmental Enrichment Plans

**DOI:** 10.1101/341206

**Authors:** Mackenzie B. Dutton, Peter J. Pierre, Jeremy D. Bailoo, Emily Warkins, George F. Michel, Allyson J. Bennett

**Author notes:** Corresponding Author: Allyson J. Bennett, Department of Psychology,University of Wisconsin-Madison, 22 North Charter St., Madison, WI 53715 USA.

## Abstract

The housing and care of captive nonhuman primates (NHP) typically meets federal regulations and standards as well as guidelines by private accreditation organizations. There is, however, a gap between such policy, common practices, and the findings of a large empirical research literature on the effects of environmental enrichment (EE), particularly with respect to the degree to which different enrichment strategies lead to a demonstrable improvement of the animal’s psychological wellbeing. Assessment tools to guide decisions about selection and refinement of EE practices are largely missing and our companion paper offers a theoretically grounded qualitative approach to the categorization and assessment of sensory, motor, and cognitive (SMC) EE strategies. Here, we propose and illustrate a model for quantitative assessment of enrichment practices using a sample of research facility, zoo, and sanctuary NHP environmental enrichment plans (EEP). Our scoring technique provides a means for comparing the efficacy of different strategies across facilities and allows for the selection of priority areas for improvement. Overall, our assessment tool provides a framework that has several advantages. It is inherently flexible. It can be tailored to fit a range of species. It can readily be adapted to accommodate new evidence about a specific EE strategy, or new EE strategies, or both. Because a scientifically valid evidence-based framework drives priority, our method is readily adaptable to different types of facilities and is more likely to lead to longer-term benefits, both in terms of the enhancement of psychological wellbeing of captive NHP, and with respect to the judicious use of limited resources.

**Acronyms:**

NHP: nonhuman primates
EE: environmental enrichment
EEP: environmental enrichment plans
SMC: sensory motor cognitive
SSIB: somatic self-injurious behavior
NSSIB: non-somatic self-injurious behavior
USDA: United States Department of Agriculture
AWA: Animal Welfare Act
AZA: Association of Zoos and Aquariums
GFAS: Global Federation of Animal Sanctuaries
NRC Guide: Guide for the Care and Use of Laboratory Animals

## 1. Introduction

Environmental enrichment (EE) is a key component of care for nonhuman primates (NHP) living in a range of captive settings that include dedicated research facilities, zoos, breeding facilities, entertainment venues, and sanctuaries across the world. Under US federal law, facilities that house NHP for exhibition (i.e., zoos, entertainment), research, or breeding are required to be licensed or registered with the United States Department of Agriculture (USDA) and must comply with federal law, the Animal Welfare Act These facilities may also operate under other rules, standards, and recommendations emanating from the various requirements and accreditation policies associated with maintenance of captive animals (for review and examples c.f., Bennett & Panicker, 2016; Hau & Bayne, 2017).

In the US, the AWA, with oversight by the USDA is the umbrella covering all licensed and registered facilities housing NHP. Thus, the AWA’s requirements are the common and core standards by which to ensure that practices and policies reflect contemporary evidence to ensure animal wellbeing. The major provision of the AWA aimed at ensuring psychological welfare is the requirement that facilities “develop, document, and follow an appropriate plan for environment enhancement adequate to promote psychological wellbeing of nonhuman primates” (*Final Rules: Animal Welfare*, 1991). The AWA requirement does not, however, identify specific types of environment enhancement/enrichment that must be used or the frequency with which they are provided. Nor is there an existing *systematic*, common-use framework for quantitative assessment from which to identify, compare, or benchmark the relative value of a wide range of enrichment strategies that are currently in use across the wide range of facilities that house NHP. The absence of a systematic framework for evaluation of NHP EE in captive settings can impose challenges to evidence-based decisions, effective practices, and rational refinement efforts. Further, there is little structure from which to prioritize hypothesis-driven research that can support best practices and be applied to advance evidence-based policy.

Advancing evidence-based practices and policies for the care of animals in captive settings is a common interest and goal not only for those working with the animals, but also for policy makers and the public. In our companion paper we have proposed a theoretically-guided model for evaluation of EE for NHP living in a range of captive settings. In the current paper we describe a novel framework for evaluation of core aspects on NHP care and illustrate how it can be applied to: 1) produce a quantitative assessment of NHP EE Plans (EEP) and guide decisions for continuing refinement; 2) identify critical gaps in knowledge about the relative effectiveness of different EE strategies; and 3) highlight key priorities for empirical research that can inform advances in NHP care.

## 2. Approach to developing a framework for assessment and a measurement tool

Our approach to developing a systematic strategy for assessment of EE was stepwise, with each component described below. In brief, we began by defining a range of EE practices in common use in a wide range of facilities that house NHP. We then evaluated the published evidence about the effect of those practices on animals – either their use of EE or its effects on behavior and wellbeing. Based on this evidence, in combination with decades of evidence from psychological science (see companion paper), we created a model quantitative tool that could be used for assessment of NHP EEP that are required by federal law. To illustrate the tool’s potential use and generality we applied it in assessing NHP EEP from a sample of different types of facilities (i.e., research institutions, zoos, and sanctuaries) that house NHP.

### 2.1. Description of the EE range evaluated here

Our first step was to identify major components of current practices and strategies for EE, with a focus on sensory, motor, and cognitive environmental enrichment (SMC EE) in the non-social, non-structural domain (see companion paper for complete discussion; also see Keeling, Alford, & Bloomsmith, 1991). As defined here, the SMC EE domain includes a broad range of enrichment: provision of toys, foraging puzzles, videogames and learning tasks, radio, television, olfactory stimuli, etc. It does not include either the social domain, which is defined in terms of housing with conspecifics, or the structural domain, which includes perches, swings, climbing apparatuses, and other large structural components of NHP housing. The rationale for focusing on SMC EE is provided in detail in our theoretical model that is the subject of our companion paper.

In brief, we propose that the SMC EE domain is critical in light of the complex cognitive, sensory, and manipulative capacities of NHP. The social and structural domains of EE are far easier to assess than SMC EE. For instance, quantifying whether animals are socially-housed, the amount of space they have, or the presence of climbing structures are all simple to objectively measure. However, both social and structural features of EE can be highly variable across facility types given that variation in physical structure, economic, and geographic factors (e.g., climate and indoor/outdoor housing, space), as well as the purpose for which the animals are housed (e.g., exhibition, breeding, research). As a result, it is the SMC EE domain that is the least well addressed in both policy and the empirical literature. That gap also continues to pose challenges for assessment, refinement, and systematic research.

The major categories and primary characteristics of SMC EE evaluated here are illustrated in Figure 1. The figure also highlights factors related to EE implementation that may affect its effectiveness. Given the wide latitude in regulation, enrichment strategies vary widely across the range of facilities that house NHP. Our goal was to propose an evaluative rubric that is *inclusive* of current practices across a *wide range* of facilities and that also permits categorization and evaluation of emerging, new, or novel practices. We explicitly focused on all types of facilities that house NHP because we believe that advances in policies and practices for the care of NHP should promote equity across settings rather than uneven standards (Bennett, 2015; Bennett & Panicker, 2016). Thus, to identify a broad sample representing the range of specific EE strategies currently in use in the US, we reviewed sources that could capture common SMC EE strategies. Those sources include EEP from different types of facilities (zoo, sanctuary, research facilities. We also analyzed standards and guidelines from accreditation organizations that represent a range of facility types and that are considerably more detailed than the AWA, including standards and best practice recommendations from the American Zoological Association (AZA), the Global Federation of Animal Sanctuaries (GFAS) and the Guide for the Care and Use of Laboratory Animals (NRC Guide). Finally, we reviewed published surveys on NHP EE practices (Baker, 2016; Baker, Weed, Crockett, & Bloomsmith, 2007) and other relevant publications appearing within the last 15 years. These included reviews, chapters, and articles (Bloomsmith, Perlman, Hutchinson, & Sharpless, 2018; Keeling et al., 1991; Liss, Litwak, Tilford, & Reinhardt, 2015; Maple, 2007; Reinhardt & Reinhardt, 2008; Schapiro, 2017; USDA Animal Welfare Information Center, 2005; Winnicker et al., 2013; Wolfle, 2005).

**Figure 1.**
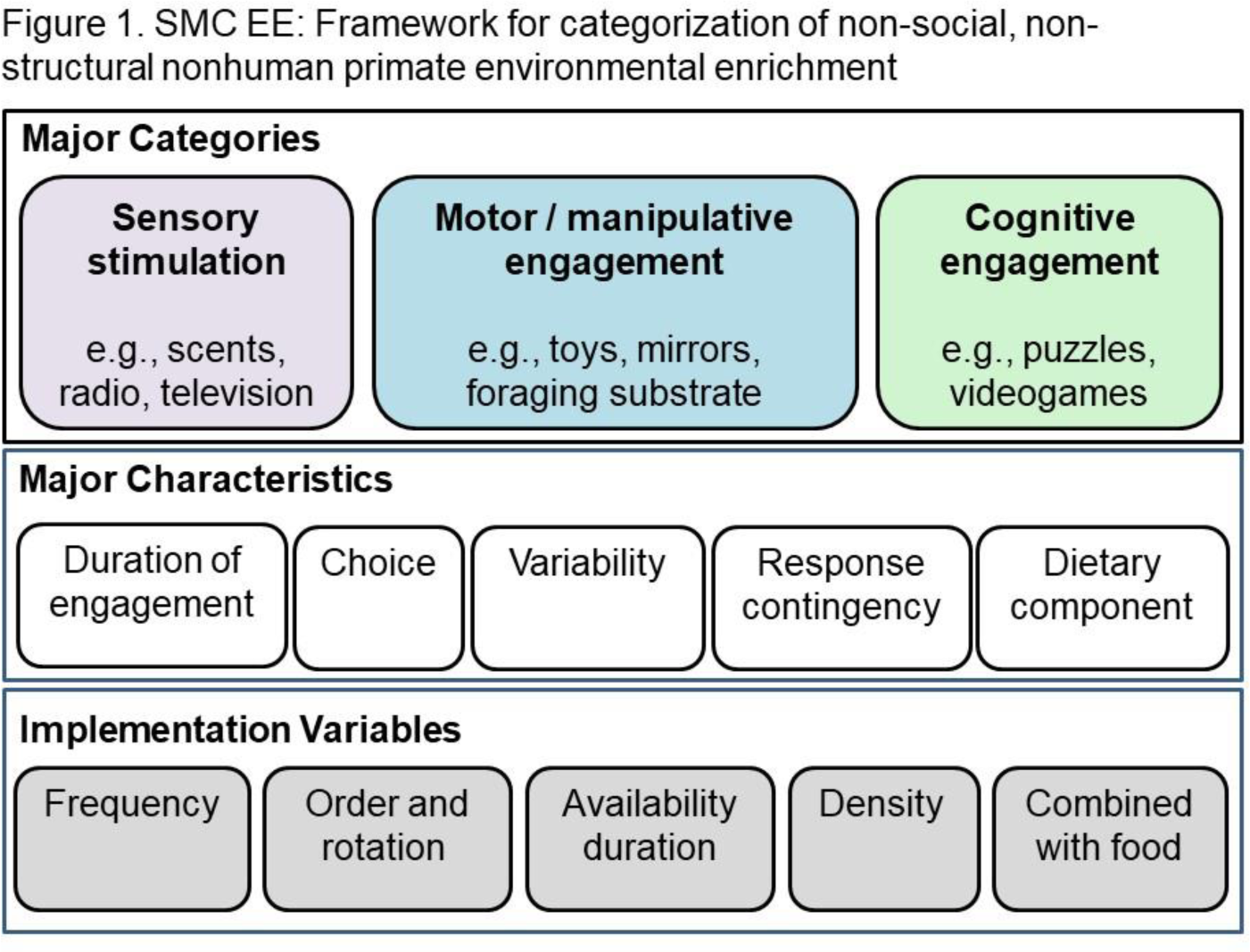
Overview of the major categories, characteristics, and implementation factors evaluated in the assessment method for non-social, non-structural/space elements of nonhuman primate environmental enrichment plans.

Our rationale for including a wide range of sources is that they provide a broader view of practices, beyond those reflected in the empirical literature. Indeed, one of the hallmarks of enrichment practices may be the contrast between the ubiquity of a practice and the limited empirical evidence to support its effectiveness. The enrichment strategies identified in our review were diverse. They included provision of manipulanda (i.e., Kong Toys™ or other manipulable devices, dolls, blankets, brushes); mirrors; television; radio; devices to promote foraging (i.e., puzzles that contain food); substrate to promote foraging (i.e., wood shavings or hay into which seeds or food could be dispersed); computerized games (i.e., games or learning tasks via video joystick or touchscreen devices); various olfactory stimulation (i.e., potpourri, scented shampoo). In addition, most plans included provision of fresh fruits, vegetables, seeds, “popsicles” or ice treats, and other foods supplemental to the chow that provides primary nutrition.

### 2.2. Evaluation of evidence for each EE strategy: Literature review

The qualitative categorization proposed in our companion paper can be useful for initial assessment and for identifying gaps and areas for prioritization of EE efforts. However, quantitative assessment, including a method to compute overall scores and, in turn, benchmark an EEP, requires a systematic approach to weighting the different strategies. Thus, the ideal scoring system is constructed with weights (or scores) assigned to reflect the relative benefit for animals’ psychological wellbeing, species-typical behaviors, and cognitive stimulation, as demonstrated by empirical study.

Following categorization of each of the common enrichment strategies identified in our survey of sources we then conducted a focused literature review. The purpose of the literature review was to illustrate an approach to identifying the breadth and strength of representative empirical evidence supporting the use of the various enrichment strategies (see Table 1 below). On the basis of the strength of both the theoretical rationale as well as empirical evidence we then assigned a proposed weight to each category of enrichment (see Table 2 and further discussion below). Ultimately the weights would be assigned based in the results of meta-analysis. To conduct a meaningful meta-analysis, however, requires a sufficient number of studies, subjects, and observations. As it stands, the number of available studies and detail in the published reports is largely inadequate to provide sufficient empirical evidence for many EE strategies commonly in practice. Thus, a meta-analysis was not conducted here. Nonetheless, the model, literature review, and analysis we present provides a structure that goes beyond identifying critical gaps in evidence and instead offers a strategic, hypothesis-driven approach to analyze and address these gaps. We summarize the gaps and approach in our conclusion section and propose priorities for future research (see below).

**Table 1.** Summary of focused literature review

| Enrichment Strategy | Studies Reviewed | Species Studied | Assessment Duration | Primary Dependent Variables | General Conclusions |
| --- | --- | --- | --- | --- | --- |
| **Computer Apparatus**<sup>1-15</sup> | 15 | Baboons; rhesus macaques; bonnet macaque; chimpanzees; orangutans; gorillas | 10 days - 7.5 months | Observational behavioral data; engagement; number of trials; cortisol (salivary) | The majority of studies found that the nonhuman primates initiated engagement quickly, increased number of trials with repeated exposure, and sustained use for prolonged periods of time without habituation. Pellet rewards were used in most studies and proven to be effective. The efficacy of video rewards versus pellets is inconsistent. Performance improved when the primates were able to choose the task. Behavioral impacts include a reduction in resting, self, and cage-directed behaviors and an increase in locomotion, social and task-directed behaviors. One study also found a reduction in salivary cortisol levels when computer system were available. |
| **Food Puzzle**<sup>16-26</sup> | 11 | Rhesus macaques; chimpanzees; stump tail macaque; pig tail macaque; common marmoset | 6 days - 10 weeks | Observational behavioral data; engagement | Food puzzles promote species normative activity budgets by requiring greater manipulation and increasing time spent foraging. The varying complexity of puzzle feeders correlates with differing degrees of fine motor skill use, cognitive engagement, and foraging time. The length of sustained engagement elicited by puzzle feeders is inconsistent. A reduction in abnormal/agonistic behaviors has been observed when puzzle feeders are available. |
| **Foraging Substrate**<sup>27-31</sup> | 5 | Rhesus macaques; stump tail macaques; chimpanzee; moustached guenons; vervets; ring-tailed lemurs; squirrel monkeys; white-headed capuchins; black-capped capuchins; red-bellied tamarins; common marmosets | 1 week - 39 weeks | Observational behavioral data; cortisol (urinary) | Foraging substrate was consistently found to promote species typical behaviors including increased foraging time, locomotion, and social behavior in a numerous species. Effects were sustained with prolonged exposure. The substrate material mattered and was significant for behavioral effects. No significant changes in cortisol levels observed. |
| **Typical Foraging Device**<sup>32-41</sup> | 10 | Rhesus macaques; cynomolgus macaques; squirrel monkeys; callitrichids; marmosets; cotton top tamarins | 2 days - 6 months | Observational behavioral data | Increases in foraging time were observed across the studies, often with associated increases in activity levels. Device type impacted the degree of initial and sustained engagement. Decreases in abnormal behavior were also observed in multiple studies. |
| **Manipulanda**<sup>42-47</sup> | 6 | Baboons; pig tail macaques; rhesus macaques; vervets; chimpanzees | 10 days - 4 months | Observational behavioral data; activity levels | Manipulanda was found to be effective in eliciting engagement and reducing abnormal behavior. However, the findings were inconsistent and engagement was found to vary by individual, throughout the day, and over time. |
| **Mirrors**<sup>48-51</sup> | 4 | Rhesus macaques; vervets; capuchins | 6 days - 30 days | Observational behavioral data; engagement | Mirrors were shown to be an effective, time-consuming enrichment device in eliciting social behavior and manipulation. Size and novelty of mirror position impacted the level of engagement. |
| **Television**<sup>52-55</sup> | 4 | Rhesus macaques; pigtail macaque | 5 weeks - 1 year | Observational behavioral data; engagement; lever pressing to turn device on/off | No clear behavioral effects identified. Engagement or looking behavior was increased when videos were shown on the screen versus blank screens, but with rapid habituation and decline in attention to the screen. The provision of television was found to be a weak positive reinforcer for lever pressing. |
| **Radio**<sup>56-57</sup> | 2 | Chimpanzees; baboons | 2 weeks - 6 months | Observational behavioral data; blood pressure and heart rate | Limited evidence to demonstrate consistent behavioral effects. Reduced activity and agitation have been observed with the provision of radio. Physiological effects, such as a lowered heart rate, have also been recorded. |
| **Enriched Environment**<sup>58-68</sup> | 11 | Rhesus macaques; cynomolgus; macaques; capuchins; chimpanzees | 5 days - several years | Observational behavioral data; cortisol (plasma, fecal); blood serotonin | The use of multiple enrichment strategies decreased abnormal behaviors and increased species typical behaviors, foraging, and activity levels. Continued, long-term use of the devices was exhibited by multiple studies. Positive effects were observed in both singly and pair-housed animals. A reduction of cortisol levels compared to baseline was also recorded in multiple studies. |

**Table 2.** Scoring tool for quantitative assessment of non-social, non-structure cogntive, motor, and sensory

| Point designation | Major Categories |  |  | Major Characteristics |  |  |  |  | Total possible points (per strategy) |
| --- | --- | --- | --- | --- | --- | --- | --- | --- | --- |
|  | Cognitive engagement | Motor/ manipulative engagement | Sensory stimulation | Duration of engagement (short-sustained) | Choice vs required | Response contingent (active vs passive) | Variability | Dietary Component |  |
| Point range | 0-3 pt | 0-3 pt | 0-3 pt | 0-2 pt | 0-1 pt | 0-1 pt | na | na |  |
| Weight/multiplier | x4 | x3 | x2 | x1 | x1 | x1 | na | na |  |
| **Enrichment Strategy** |  |  |  |  |  |  |  |  |  |
| Computer-based Device | 3 | 3 | 2 | 2 | 1 | 1 | Absent/<br>present | Likely | 29 |
| Puzzles | 3 | 3 | 2 | 2 | 1 | 1 |  | Present | 29 |
| Foraging Substrate | 2 | 2 | 3 | 2 | 1 | 1 |  | Present | 24 |
| Foraging Objects | 2 | 2 | 2 | 1 | 1 | 1 |  | Present | 21 |
| Manipulanda* | 1 | 1 | 1 | 1 | 1 | 1 |  | Absent | 12 |
| Mirror | 1 | 1 | 1 | 1 | 1 | 1 |  | Absent | 12 |
| Television** | 0 | 0 | 2 | 0 | 0 | 0 |  | Absent | 4 |
| Radio** | 0 | 0 | 1 | 0 | 0 | 0 |  | Absent | 2 |
| **Total possible** | **48** | **36** | **28** | **9** | **6** | **6** |  |  | **133** |

Scale: 0-31 (**Quartile 1/Minimal**); 32-65 (**Quartile 2/Average**); 66-99 (**Quartile 3/Exceeds common practice**); 100-133 (**Quartile 4/Optimal**)
\*Manipulanda without food inserted; \*\* Television or radio without option for animal to control the device

### 2.3. Application of assessment tool

The overall goal of this paper and its companion is to offer a systematic approach to evaluation and advancement of EE in the SMC domain. The approach is guided by psychological theory relevant to consideration of wellbeing and by empirical evidence about the effects of different forms of enrichment. We offer the following caveats: The approach here is not meant to be an exhaustive catalog of studies, reports, and anecdotal observations of all variants of SMC EE. While such catalogs and comprehensive reviews have merit and utility, without further elaboration and organization into systematic frameworks they are unlikely to result in broad advances in effective policy and practice. Nor is the analysis included in this paper meant to be a systematic review of EEP in US facilities, or even an evaluative comparison and judgement of the relative strength and effectiveness of enrichment programs in different facility types. To provide an *initial illustration* of how the proposed framework could be used, we applied the assessment strategy to a set of 15 NHP EEP that we obtained by request from 8 research facilities, 5 sanctuaries, and 2 zoos. Representative results of the analysis are provided in Table 3 (see Section 5 below).

**Table 3.**
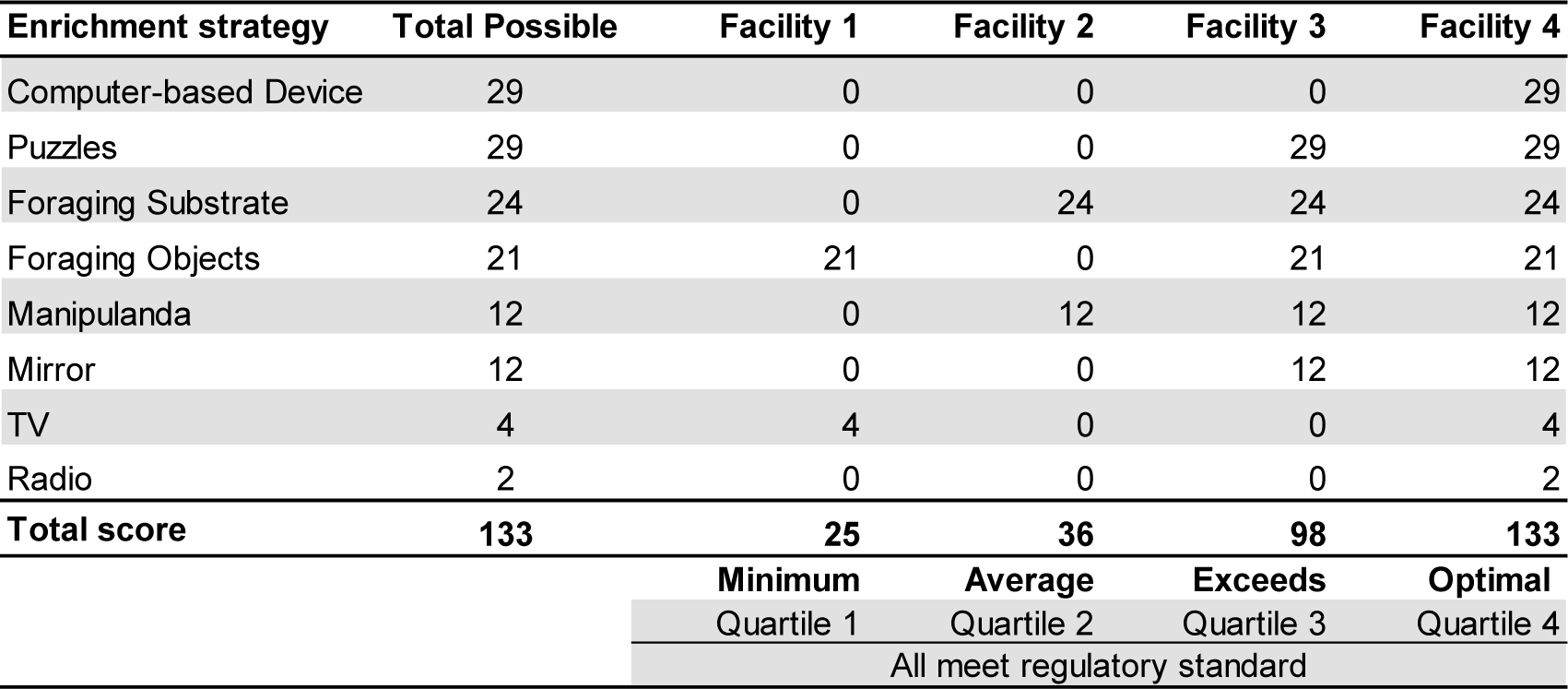
Illustration of assessment tool applied to four facility’s environmental enrichment plans (EEP)

## 3. What is the evidence for the value of different EE strategies? Rationale and description of focused literature review supporting the assessment framework

The scientific literature to support selection of EE strategies is uneven and significant gaps remain regarding the relative effectiveness of different strategies. Moreover, the situation for NHP stands in contrast to a robust and long-standing *experimental* literature focused largely on rodents (rats, mice) and on the evaluation of EE on brain, behavior, immunology, and health in animals without clinical concerns (Benefiel, Dong, & Greenough, 2005; Bennett, Diamond, Krech, & Rosenzweig, 1964; Fischer, 2016; Frasca, Tomaszczyk, McFadyen, & Green, 2013; Hebb, 1949; for review c.f., Renner & Rosenzweig, 1987; Simpson & Kelly, 2011). These rigorous experimental studies investigated such causal relationships through random assignment of animals to groups without EE and with variations of EE.

In contrast to rodents, NHP experimental studies examining the causal effect of social and physical environmental variables reached a peak in the period around 1950-1980; there were no parallel contemporary efforts in NHP. This is likely, in part, because the use of a no (or low) enrichment control group would be ethically challenged and require exemption from federal and community standards. The absence of rigorous experimental studies aimed at determining the causal effects of specific EE strategies on brain, behavior, and physiology means that conclusions about EE are typically drawn from short-term studies that may use both weak manipulations of the independent variable (EE) as well as insensitive dependent measures (see discussion below). Nevertheless, a review of the extant literature can be used to illustrate how available data, even when limited, can be used to assess the relative effectiveness for each of the commonly-used EE strategies.

### 3.1. Measuring the effects of EE on animal behavior and wellbeing

Decades of research have investigated the effects of EE on behavior and outcome measures thought to be relevant as markers of wellbeing in NHP. Part of the difficulty in defining the benefit and relevance of EE, and thus the strength of the EEP, lies with the lack of consensus of an operational definition of wellbeing. One definition of wellbeing refers to “the state of an animal as regards its attempts to cope with its environment” (Broom, 1986). Whether an animal is successfully “coping” with its environment can be expressed in different ways and has led to different approaches to the assessment of animal welfare that overlap and complement each other. These include naturalistic, function-based, and feelings-based approaches.

The naturalistic approach refers to the animals’ behavioral “integrity” and, for example, assesses to what extent animals can express “species-typical” behavior (e.g., foraging). The function-based approach refers to the animals’ normal biological functions and assesses to what extent the conditions under which animals are kept compromise these functions (e.g., stress physiology). The feelings-based approach refers to the animals’ affective or mood states. As feelings involve subjective experiences, they can only be inferred from measures of affective states that are not confounded by the animals’ activity and emotional arousal, such as cognitive bias (Harding, Paul, & Mendl, 2004).

With respect to federal regulation and policies, the AWA specifies two goals that are associated with improving NHP wellbeing: 1) promoting species typical behavior; and 2) decreasing abnormal behaviors. These two goals are subsumed under the naturalistic and function-based approaches, respectively. The implicit rationale reflected in these goals is that species-typical behaviors are viewed as positive, or reflective of a species’ natural history, including foraging behaviors, manipulation/exploration, increased activity levels and time spent feeding (Lutz & Novak, 2005). Reliable assessment tools under the feeling-based approach are lacking, and those, which show promise, such as assessment of judgment biases, are relatively new and require further validation; the feelings-based approach will not be covered in the present manuscript.

#### 3.1.1. Clinical treatment model

Research on the effects of EE has often focused on the reduction of what are termed “abnormal” behaviors, including stereotypies (i.e., rocking, body flips), self-directed behavior (i.e., digit-sucking, hair pulling), self-injurious behavior (i.e., self-biting, self-wounding) that occur in a some primates in a broad range of captive settings (for review c.f., Jacobson, Ross, & Bloomsmith, 2016; Lutz, 2014; Mason, 1991; Novak, Hamel, Ryan, Menard, & Meyer, 2017). We view this type of research as approaching evaluation of EE from a “clinical treatment” model, one that is focused on reducing behavioral symptoms that can be associated with compromised psychological and physical welfare. Among the underlying assumptions in the clinical treatment model are several for which there is little direct evidence. One is that animals without such symptoms also benefit from the same EE that is useful to “treat” symptomatic animals. Further, and related, is the assumption that treatment SMC EE strategies may be prophylactic for non-symptomatic animals. That is, “treatment EE” may reduce the likelihood or development of symptoms or full-blown wellbeing problems (i.e., self-injurious and abnormal behavior).

A robust and long-standing literature clearly points to the importance of social housing to reduce risk of abnormal and self-injurious behavior. There are few (if any) large prospective or retrospective studies in NHP that could provide evidence for the prophylactic effectiveness of SMC EE, however. Nor is it clear that such a study would be practically or logistically feasible. In part this is because implementation of SMC EE is so widely variable across facilities and across time (see above and companion paper). In rodents, however, where prospective study and manipulation of relatively homogenous housing environments and uniform standards are possible, there are a handful of studies which provide some support of the notion that SMC EE can *reduce* levels of, but not prevent, abnormal behavior (Bechard, Meagher, & Mason, 2011; DeLuca, 1997; Gross, Engel, Richter, Garner, & Würbel, 2011; Latham & Mason, 2010; Nevison, Hurst, & Barnard, 1999; Tilly, Dallaire, & Mason, 2010; Würbel, Chapman, & Rutland, 1998). It is critical to note here, that these studies are few in number, evaluated a range of abnormal behaviors (e.g., stereotypy, barbering) and SMC EE strategies and, perhaps most importantly, subsequent replications have failed to find consistent effects when similar categories of abnormal behavior are studied in relation to SMC EE strategies (Bailoo et al., 2018; Gross et al., 2011; Gross, Richter, Engel, & Würbel, 2012; Latham & Mason, 2010; J. Novak, Bailoo, Melotti, & Würbel, 2015).

The treatment model is important to serve animals that exhibit compromised wellbeing. At the same time, it is also critically important to identify what proportion of animals exhibit abnormal behavior. Estimating the prevalence of such behaviors can inform decisions about whether the clinical treatment model should be used. Moreover, it is needed to evaluate whether abnormal behavior is a robust outcome measure or, conversely, an insensitive dependent variable, in evaluation of EE strategies. In other words, if the prevalence of abnormal behavior is low at baseline it is unlikely to provide a useful measure that discriminates effective EE strategies. Further, we might consider the implications if the prevalence of abnormal behavior is low and whether the assumptions outlined above— that SMC EE is prophylactic and that it equally benefits animals with and without compromised wellbeing— are wrong. If this is the case, then studies focused on abnormal behavior and the treatment model will fall short of advancing knowledge about EE that can best benefit the larger population of NHP that includes many animals without significant abnormal behavior.

#### 3.1.2. Prevalence of abnormal behavior

Estimating the prevalence of abnormal behaviors that are indicative of compromised wellbeing in the NHP population in captivity is difficult. Moreover, identifying and comparing prevalence across species, age groups, facility types, housing conditions, and other factors is even more difficult. Full review is beyond the scope of this paper. Over the past 50+ years several research groups have published studies reporting on either prevalence estimates or analysis of factors associated with risk of NHP developing abnormal behavior (e.g., for review and additional references see Lutz, 2014; M. A. Novak et al., 2017). In reports published after 2000, estimates of prevalence of *abnormal behavior* are wide-ranging. For instance, a sampling of reports on the percentage of animals exhibiting abnormal behavior include: 13% of prosimian primates in a US zoo (Tarou, Bloomsmith, & Maple, 2005); 48-64% of chimpanzees in 26 accredited US zoos (Jacobson et al., 2016); 89% of singly-housed rhesus macaques in a former US research facility (Lutz, Well, & Novak, 2003); and 90% of singly-housed cynomolgus macaques in a Chinese research facility (Camus, Blois-Heulin, Li, Hausberger, & Bezard, 2013).

There is high variation in categorization and coding schemes for NHP abnormal behavior. Here we will distinguish between *somatic self-injurious behavior* (SSIB; i.e., self-wounding that is severe and requires medical treatment) and all other forms of abnormal behavior, including self-biting and mouthing that does not result in wounding (i.e., *non*-somatic self-injurious behavior; NSSIB). Estimates of prevalence of SSIB are generally much lower in both research and zoo-housed animals. For example, in laboratory-housed macaques, there are reports of 3% (Bellanca & Crockett, 2002) and 11% (Lutz et al., 2003). A study of 231 rhesus monkeys reported that over 50% exhibited what the authors termed “self-abuse,” but there was insufficient detail to yield prevalence estimates for SSIB (Rommeck, Anderson, Heagerty, Cameron, & McCowan, 2009). In a study of chimpanzees in 26 accredited US zoos, roughly 2% were reported to exhibit self-injurious behavior (Jacobson et al., 2016). Similarly, roughly 2% of 540 primates housed in 35 zoos and including prosimians, New and Old World Monkeys, and apes in British and Irish zoos were reported to engage in self-biting (Hosey & Skyner, 2007). By contrast, a study of chimpanzees living in 40 US and UK zoos reported that 20% of the 40 chimpanzees engaged in self-biting (Birkett & Newton-Fisher, 2011), though it is not clear whether this was SSIB. There is relatively little data about animals housed in sanctuaries or other settings.

What is not reflected in the published literature is how many animals and populations *do not* exhibit either NSSIB or SSIB. The great majority of reports on both are from a handful of colonies in which abnormal behavior is either a focus of study or a phenotypic measure collected as part of other research. It is possible that the prevalence estimates from those colonies may generalize to others. Alternatively, there may be low external generalizability. Some will argue that in absence of evidence, we should assume external generalizability. That is, the prevalence estimates are indicative of rates of SSIB and NSSIB in other colonies for which there is no published data. However, doing so depends on meeting the assumption that the studied colonies are representative of the larger population. It is not at all clear that the assumption is true. Further, if the assumption is not tested explicitly, the generalization remains weak. It therefore remains for future study to test whether the prevalence of SSIB is similar across colonies and settings. The current published literature is insufficient to produce strong conclusions on this point.

Interpretation and generalization of the findings on prevalence of abnormal behavior in NHP is also challenged by several historical, definitional, and methodological considerations. One clear challenge is the fact that standards of care for NHP have changed over time. Specifically, housing and care conditions, including the provision of social-housing and multi-modal EE have been the focus of attention and has changed via practices and federal policy over the past three decades (for review c.f., Novak et al., 2017; Schapiro, 2017; USDA Animal Welfare Information Center, 2005; Weed & O’Neill-Wagner, 2015). Thus, the rearing and care conditions for NHP in the 2000s are unquestionably different than those for animals prior to the 1996 changes in federal law.

Given the very reason for the policy changes was to improve animal wellbeing, it seems probable that animals reared in earlier housing, care, and EE conditions may exhibit different behavior than those reared in contemporary conditions. However, in viewing the entirety of the literature on abnormal behavior in NHP it is clear that most of the citations and findings are from studies in the 1950s through the 1990s, studies of animals housed under very different conditions. Even in studies published after 2000, the animals surveyed for incidence of abnormal behaviors were often born and reared under pre-1996 conditions. There are very few reports on the prevalence of self-injurious, self-directed, and abnormal behaviors in animals who were born and reared entirely under contemporary standards for housing, care, and EE. An additional complication is the fact that there are very different NHP populations with respect to experiential history. There are populations who were born into multiple generation domestic breeding colonies, then reared and housed in similar conditions throughout their lives. There are also populations that were born and reared in one setting and then transferred to very different housing conditions. For instance, some animals imported for zoos, breeding, or research may have lived in facilities with large outdoor housing and then been transferred to smaller indoor housing. In the case of zoos and sanctuaries, some animals may have lived in private homes and then been transferred to social groups and facility housing (Jacobson et al., 2016).

Another final factor to disentangle is the effect of cross-generational and social influences on the expression of behavior. Hook and her colleagues (2002) have provided evidence from a study of 8 chimpanzee groups and 21 rhesus macaque social groups that: “Variance in the expression of abnormal behaviors across groups of chimpanzees and rhesus monkeys suggests that social learning processes are involved in the propagation of these behaviors and thus, the study of abnormal behaviors may be a suitable method for further examining the question of culture in non-human primates.” In the same vein, others have concluded that a high prevalence of a specific NSSIB behavior indicates that it is social transmitted and should not be considered abnormal (e.g., coprophagy in zoo chimpanzees; Jacobson et al., 2016).

Overall, analysis of cohort, social group, age, and experiential history are all needed to better understand the prevalence of abnormal behavior and, in turn, address *whether* it is a robust and meaningful outcome measure for the broad range of studies that evaluate EE. Further, consideration of the complex factors that may influence the incidence of abnormal behavior are also needed to produce accurate prevalence estimates.

#### 3.1.3. Definitional and methodological issues in measurement of abnormal behavior

Other challenges to interpreting the extant literature on prevalence of abnormal behaviors arise from intertwined definitional and methodological issues. In brief, some behavioral symptoms that are indicative of compromised wellbeing can be readily quantified. For example, SSIB leads to flesh wounds that require medical treatment, as such, it is readily and objectively measurable and available in medical records. Thus, a large-scale study of the incidence of SSIB is feasible. Further, there is little disagreement that SSIB is a clear indicator of compromised wellbeing.

By contrast, there is disagreement and a lack of evidence that some of the behaviors typically designated as markers of compromised wellbeing are valid indicators of diminished wellbeing. Primary among these are pacing and mild-moderate persistent alopecia. Both may occur in a large proportion of animals, but there is no clear evidence that either significantly compromises the animals’ health or their engagement in typical behaviors for their species, age group, housing, etc. (Novak et al., 2017). In fact, Poirier and Batson (2017) have recently provided a detailed critical evaluation that argues for the urgency of determining whether pacing is, in fact, an indicator of wellbeing. They observe:

> “Our review highlights the current lack of understanding of the causal factors underlying pacing behavior. According to current knowledge, the welfare of pacing macaques could be either better, worse or equivalent to that of nonpacing individuals. It is also unclear whether pacing results from brain abnormalities. Since rhesus macaques are widely used as a model of healthy humans in neuroscience research, determining if pacing behavior reflects an abnormal brain and/or poor welfare is urgent”

Beyond definitional issues are those that are compounded by the methods commonly used for colony-wide or large population surveys. The majority of systematic, published studies of abnormal behavior come from studies of animals in research settings. The methods often involve very short (i.e., 5 minutes) and infrequent (i.e., once or twice a year) observations by a human observer. The data collected may be a simple yes or no for each of a range of behaviors. One reason for the brief evaluation is practicality. These studies are typically part of wellness assessments or large-scale research assessments for very large colonies of animals. Thus, apart from self-wounding and alopecia, the typical measurements of abnormal behavior are based in a very brief time window. As a result, the measurement may miss behaviors entirely, or may yield an inflated estimate of their occurrence over a typical day, week, or month. That is, an abnormal behavior exhibited one time in two 5-min observation periods might be a reliable marker of what occurs frequently, but it also may not. The unvalidated assumption in this approach is that the behavior is relatively stable and its occurrence under small windows of observation generalizes to non-observed times.

Overall, because of the issues related with measurement, validation, and ethological relevance, it is difficult to draw firm conclusions with respect to true prevalence of abnormal behavior of captive NHP. Some studies report high levels while other do not. Moreover, the heterogeneity in types of abnormal behavior, as well as the heterogeneity of breeding, rearing, and housing conditions of NHPs, make the drawing of *general* conclusions about the etiology and treatment of abnormal behavior difficult. Whether EE that is used as treatment for animals with symptoms is also beneficial to other animals is similarly understudied. Arguably, in absence of strong evidence that the benefit of SMC EE generalizes from symptomatic to other animals, research on EE should clearly differentiate between the treatment model and evaluation of EE effects on the larger population of animals without symptoms such as SSIB, stereotypies, and other “abnormal behavior.” Further, given the possibility that the prevalence of such behaviors is low, its use as an outcome measure may therefore create a “floor effect” problem, or be an insensitive dependent variable with respect to adequate assessment of the effects of EE.

#### 3.1.4. Objective measures: Engagement and use of EE by animals

A different, and complementary, approach to measuring the effectiveness of EE is to record usage, that is whether, to what extent, how long, or how frequently animals choose to interact with it. Such measures do not necessarily provide evidence of benefit in terms of wellbeing. However, absence of interaction or interest from the animal, does provide a readily quantifiable marker that likely reflect ineffectiveness of the EE. In fact, several groups have argued that measurement of animals’ engagement should be the common initial assessment for any EE strategy (Bennett et al., 2014; Lutz & Novak, 2005; Schapiro & Lambeth, 2007). Thus, in this review we include measures of sustained engagement as key information for assessment (see also Section 5.2 below).

The concept of engagement is reflected in much of the literature about EE and is a frequent outcome measure. It is also included in broad categorization schemes that focus on behavioral and sensory affordances that EE offers the animal. For example, Lutz and Novak (2005) describe categorization of inanimate enrichment in terms of: “… those that require some physical activity on the part of the animal (active enrichment) and those that provide only passive kinds of stimulation… This distinction is somewhat arbitrary inasmuch as passive forms of enrichment can be converted to active forms of enrichment if the animal can control the onset or offset of exposure. Furthermore, active forms of enrichment may provide only passive stimulation if the animal does not use them.” (page 179-80). In this categorization “active” EE practices (i.e., those that permit some level of engagement or activity by the animal) would include provision of foraging devices, manipulable objects, fruits, and vegetables. By contrast, passive enrichment might include exposure to television, radio, or olfactory stimuli. Engagement with active EE can be measured by manual, pedal, oral, or other direct contact. Engagement with passive stimuli can also be measured. For example, by recording visual attention directed at a television, contact or proximity to a localized olfactory stimulus, or through providing the animal the opportunity to turn the stimuli on or off.

#### 3.1.5. Wellbeing and outcome measure summary

Overall, the current literature on the relative effectiveness of various EE strategies poses some challenges to advancing evidence-based practices and policies. Among them is the need for additional research to refine and operationalize the concept of wellbeing in NHP. While behaviors typically associated with compromised health – self-injurious behavior, stereotypies, excessive aggression (i.e., those often referred to collectively as abnormal behavior) – are relatively well-defined, there remains variation in methods of measurement and recording. Consensus on the definition and measurement of wellbeing remains elusive. Beyond the general agreement that presence of abnormal behavior indicates a lack of wellbeing, other indicators may include: physical condition (including coat), activity levels, and stress hormone levels typical for animals of the same the age, sex, species, and housing; engagement in species-typical behavior (e.g., foraging, play, eating).

Continued refinement and normative values for all these potential markers of wellbeing will facilitate advances in measurement that allow for quantification and comparison that are at the heart of evidence-based assessment of the effect of different EE, care, and housing practices. For the purposes of this review and development of the assessment framework we included all outcome measures without attempting to impose subjective relative value with respect to wellbeing. In other words, we treated, for example, reports of increased activity and reduction of stereotypies equivalently. Clearly, however, the tool proposed here could readily be adapted with new evidence supporting differential categorization and weighting. It could also be adapted to create individualized treatment plans tailored to prioritize EE strategies with documented benefit for particular abnormal behaviors.

### 3.2. Specific EE practices included in the assessment tool

In this paper we focus on a subset of EE strategies in the SMC domain that meet three criteria. First, they are in broad use across types of facilities, as indicated by their mention in the EE standards for research facilities (Guide), zoos (AZA), and sanctuaries (GFAS). Second, they appear to be in common use (e.g., see Baker, 2016; Baker et al., 2007). Third, they are identified in our companion paper as candidates based on broad theoretical principles and evidence from psychological science. The strategies that were included are: computer-based, puzzle feeders, foraging substrates, foraging objects, manipulanda (toys), mirrors, television, radio, and multiple types (see Table 1).

Some EE strategies that are commonly mentioned and in use are not included in our review due to a near absence of studies investigating their efficacy. For example, supplemental feeding, or giving animals a range of foods and treats, without the accompaniment of a puzzle, foraging device, or other means of promoting manipulation, is not included. Supplemental feeding can clearly offer sensory stimulation and may have great value, however, there is little direct study of its effect. Similarly, provision of novel scents and olfactory stimuli may have value, but there is very little direct study or evidence from which to evaluate its impact.

## 4. Evidence related to SMC EE strategies included in assessment tool: Literature review

As described above, a comprehensive review of each study of the EE strategies included in our assessment model is beyond the scope of the current paper and is provided elsewhere (see references above). Here we provide a synopsis of studies within the peer-reviewed literature to identify the primary findings for each of the EE strategies within the SMC domain. Below we summarize the rationale for the use of each strategy and the current evidence for its effectiveness with respect to either the animal’s wellbeing, engagement with the strategy, or both. Table 1 provides a synopsis of the findings.

### 4.1. Computer-based strategies

In these EE strategies, a computer-based device is employed in which software controls the presentation of stimuli, records the “active response” of the animal and signals resultant response contingencies (i.e. correct, error and reward presentation). The active response requires the animals to engage through touchscreen, joystick or another controller. Computer-based devices often are used to measure complex cognitive functions, such as interactive memory, recognition and problem solving.

Computer-based strategies for EE have been used in a variety of species (e.g. rhesus macaques (Bennett, Perkins, Tenpas, Reinebach, & Pierre, 2016; Washburn, Hopkins, & Rumbaugh, 1991; Washburn & Rumbaugh, 1992), bonnet macaques (Andrews, 1994; Andrews & Rosenblum, 1993, 1994a, 1994b), baboons (Fagot & Bonté, 2010; Fagot, Gullstrand, Kemp, Defilles, & Mekaouche, 2014; Fagot & Paleressompoulle, 2009), orangutans, chimpanzees, and gorillas (Calapai et al., 2017; Mallavarapu, Bloomsmith, Kuhar, & Maple, 2013; Perdue, Clay, Gaalema, Maple, & Stoinski, 2012; Platt & Novak, 1997; Wagner, Hopper, & Ross, 2016). The literature evaluating the response of NHP to computer-based EE provides evidence of effects on a range of outcome measures, including sustained engagement, task performance, behavior, and cortisol levels.

Several studies demonstrate that NHP will engage with computer-based games for extended periods of time with no evidence of habituation or reduced use over repeated presentation. For example, in one study, a computer-based joystick apparatus proved to be more effective in maintaining sustained engagement for 11 rhesus macaques compared to commonly-used foraging device – during five-hour testing sessions, all animals averaged over 1,000 trials (Bennett et al., 2016). Similar levels of engagement – over 1 million trials across 85 days – were reported for baboons housed in large social groups, with approximately 75 of all individuals in the group actively participating (Fagot & Bonté, 2010; Fagot & Paleressompoulle, 2009). Other studies have examined how factors of control and different types of rewards affect performance levels on computers. For example, rhesus monkeys were shown to have higher performance levels if they were given a choice of tasks to select from as opposed to being assigned a task (Washburn et al., 1991). The use of food pellets is a typical reward provided when the computer tasks are completed correctly. Monkeys also exhibited sustained engagement when non-food rewards, such as video clips, were used as response contingencies (Andrews & Rosenblum, 1993, 1994a), thus demonstrating, a food motivator is not required to maintain engagement with computer-based devices.

Evidence of beneficial effects of video-tasks is apparent from empirical studies that demonstrated decreases in cortisol, cage-directed, self-directed and stereotypical behaviors (for review, c.f., Bennett et al., 2016; Fagot et al., 2014; Perdue, Beran, & Washburn, 2017; Washburn & Rumbaugh, 1992). When the computer-based devices were available rhesus macaques spent 40% of their time on task related behaviors – compared to the 20% of time spent on device directed behavior observed with other enrichment devices. The monkeys also showed a reduction in self-directed, cage-directed and stereotypical behaviors when the computer-based devices were available (Washburn & Rumbaugh, 1992). On average, baboons showed an increase in activity, social-behaviors, and task-directed behaviors when allowed access to the computer-based games. They also exhibited decreases in salivary cortisol, suggesting these changes in behaviors do indeed correlate with lower stress levels and consequently benefits to health (Fagot et al., 2014).

Together, the extant literature over the past 25 years provides strong empirical support for the effectiveness of computer-based devices as an EE strategy. Additionally, computer-based tasks have inherent flexibility and capacity for use as enrichment in a wide-range of relevant characteristics, including cognitive engagement (learning, memory, attention), motor manipulation, sensory stimulation, sustained engagement, choice, and response contingency. Further, computer-based tasks can provide variability within each of these characteristics. For instance, the complexity of tasks and visual or auditory stimuli can readily be varied. Finally, computer-based enrichment is uniquely well-suited to evaluate and demonstrate clear evidence of sustained engagement because it allows for automated and continuous measurement of the animals’ use, or engagement (Bennett et al., 2016; Fagot & Bonté, 2010; Fagot & Paleressompoulle, 2009). In other words, whereas traditional puzzles or toys typically require a human observer to record the amount of time animals contact or engage with the object, a feature of computer-based tasks is continuous and detailed recording of animals’ interaction with the device.

### 4.2. Puzzle strategies

Puzzle enrichment is defined here in the terms of the provision of a device that requires a series (2 or more) of coordinated movements to solve. Often, food is used as a reward for completing the puzzle and here we refer to those as *food puzzles*. Inclusion of a series of manipulation in the definition is a straightforward way to differentiate between food puzzles and other foraging objects. While foraging objects allow supplemental food to be obtained via rote simple actions that occur quickly and without substantial engagement, food puzzles require some complexity of investigation and manipulation. Due to the lack of literature on non-food puzzles as an EE strategy, and their absence in common practices, we have focused on food puzzles here.

Puzzle feeders are a common EE strategy and their effectiveness has been assessed in a broad range of NHP, including chimpanzees (Bloomstrand, Riddle, Alford, & Maple, 1986; Brent & Eichberg, 1991; Celli, Tomonaga, Udono, Teramoto, & Nagano, 2003), rhesus macaques (Bloom & Cook, 1989; Harlow, 1950; Reinhardt, 1993b), pigtail macaques (Murchison & Nolte, 1992), stumptail macaques (Reinhardt, 1993a), and common marmosets (de Rosa, Vitale, & Puopolo, 2003; Roberts, Roytburd, & Newman, 1999). In general, these studies have examined changes in agonistic and abnormal behavior, general activity levels and foraging times, and duration of interaction with the device.

As an enrichment strategy, puzzle feeders enhance species-typical behaviors such as time spent foraging by increasing the complexity of actions required for the animal to extract the food contained within the device (Bloom & Cook, 1989; Bloomstrand et al., 1986; Murchison & Nolte, 1992). In chimpanzees housed in pairs or in social groups, sustained engagement for over an hour, reduced agonistic/abnormal behaviors (i.e., biting, hair pulling, coprophagy, feces spreading, repetitive behaviors, self-slapping), and increased foraging behaviors have been reported when puzzle feeders are provided (Bloomsmith, Alford, & Maple, 1988; Bloomstrand et al., 1986; Celli et al., 2003). Similarly, marmosets demonstrated decreased pacing and inactivity – behaviors deemed undesirable by these authors – and increase foraging time with increased environmental complexity provided via wooden gum and puzzle feeders (de Rosa et al., 2003; Roberts et al., 1999).

Puzzle feeders can vary in complexity, and consequently elicit different degrees of cognitive engagement and motor manipulation. As anticipated, an increase in puzzle complexity was found to correlate with greater manipulation and foraging time in rhesus macaques (Bloom & Cook, 1989). The extent of sustained engagement, and conversely habituation, are unclear. Brent & Eichberg (1991) found that chimpanzees spent a consistent amount of time manipulating puzzle-boards across trials. In contrast, Celli and colleagues (2003) found that chimpanzees decreased manipulation from 36.6% to 8& over three observation sessions. Other studies found a peak in manipulation initially (Roberts et al., 1999) and over the first hour of access to the device (Murchison & Nolte, 1992). Individual differences, study duration, and the impact of device complexity on habituation and manipulation are likely contributing factors to the inconsistent findings regarding sustained engagement with food puzzles.

Overall, there is strong empirical support for puzzle feeders as an effective form of EE; they provide cognitive, sensory, and manipulative stimulation and have been proven successful in a wide range of primate species. The increased complexity of obtaining food objects promotes species normative activity budgets, foraging behaviors, such as tool use, and sustains engagement for prolonged periods. While most studies focused on usage and engagement, reductions in abnormal behaviors have also been reported (Bloomsmith et al., 1988; Roberts et al., 1999).

### 4.3. Foraging substrate strategies

In addition to puzzle feeders (i.e., discrete devices), EE aimed at increasing foraging behavior can also be provided through use of dispersed substrate, or a textured medium (e.g., wood chips, wood wool, paper, shavings, etc.), within an animal’s enclosure within which food may be distributed. The number of published studies that report on the effects of foraging substrate as an EE strategy is not particularly large, however, the evidence of its effectiveness is consistent. Foraging substrates have been studied in several species, including chimpanzees (Baker, 1997), capuchin monkeys (Ludes-Fraulob & Anderson, 1998; Ludes & Anderson, 1996), rhesus macaques (Byrne & Suomi, 1991), stumptail macaques, guenons, ring-tailed lemurs, vervets, red-bellied tamarins, squirrel monkeys, and common marmosets (Chamove, Anderson, Morgan-Jones, & Jones, 1982). The literature includes reports on both the amount of manipulation of the foraging substrate as well as changes in the animals’ abnormal behaviors, activity levels, and social interactions. In one study involving 13 chimpanzees housed in pairs or trios, the addition of straw and foraging material decreased abnormal behaviors possibly by increasing foraging behavior (Baker, 1997). Other studies found similar results across a range of NHP, with increases in foraging time and overall activity levels in the presence of foraging substrate (Byrne & Suomi, 1991; Chamove et al., 1982; Ludes-Fraulob & Anderson, 1998). Furthermore, the type of substrate was found to be a significant variable in the resultant behavioral impacts (Chamove et al., 1982; Ludes & Anderson, 1996).

Together the literature provides evidence that the use of foraging substrate can produce benefits to animal wellbeing and is likely an effective EE strategy. The benefits are multifaceted. Soft bedding provides a natural substrate to bare floors, is associated with increases in activity and species typical behaviors, and can lead to reductions in abnormal behavior, aggression, and over grooming (for review and additional references c.f., Bennett, Corcoran, Hardy, Miller, & Pierre, 2010).

### 4.4. Foraging object strategies

A third category of foraging enrichment is the use of a discrete object (i.e., separable from the caging) that is designed to elicit foraging behavior through relatively simple manipulative actions required to extract food. Typical simple foraging objects are balls, boxes, or PVC tubes that have holes and are held by one hand (or foot) while the food items placed within the device are removed. The devices are categorized separately from puzzle feeders because they require simple manipulation and offer little problem-solving challenge.

Foraging devices are widespread in use and their EE value has been studied in a range of species that includes rhesus macaques (Bayne et al., 1991, 1992; Reinhardt, 1993b), cynomolgus macaques (Bennett et al., 2014; Holmes, Riley, Juneau, Pyne, & Hofing, 1995), squirrel monkeys (Fekete, Norcross, & Newman, 2000; Spring, Clifford, & Tomkol, 1997), and callitrichids (Chamove, 2005; Sha, Ismail, Marlena, & Lee, 2016; Voelkl, Huber, & Dungl, 2001).

Observational data and behavioral effects were evaluated for this EE strategy, including: amount of foraging time and duration of interaction with the device, as well as changes in abnormal behavior, grooming behavior, and activity level. Significant increases in foraging time were observed with the provision of foraging objects to single- and group-housed squirrel monkeys, marmosets, and two species of macaques (Bennett et al., 2014; Chamove, 2005; Fekete et al., 2000; Holmes et al., 1995; Voelkl et al., 2001). Similarly, when a typical food box, or hopper, that required only simple reaching to retrieve food was replaced by one providing food through wire mesh – complicating food extraction – foraging time increased as would be predicted with increased complexity. In pair-housed rhesus macaques, this simple increased manipulation requirement increased foraging time 141-fold (Reinhardt, 1993b). Provision of artificial turf boards and polyvinyl chloride (PVC) pipe devices resulted in a decrease in stereotypical behavior in singly-housed rhesus and cynomolgus macaques (Bayne et al., 1991, 1992; Holmes et al., 1995). Comparable behavior effects were observed in cotton top tamarins and Goeldi’s monkeys, which showed an increase in species typical behaviors (i.e., foraging) and activity budgets (Sha et al., 2016).

It is important to note that some devices are more effective at maintaining animals’ engagement with the device over a period. Unsurprisingly, devices that pose a manipulative challenge for the extractions of small food items (e.g., such as a PVC pipe with small holes drilled in it) sustain engagement longer than simply providing food items in an open treat dispenser (Bennett et al., 2014). Overall, typical foraging devices that encourage manipulation and engagement from captive NHP can lead to positive behavioral changes and likely are an effective form of EE (for additional review and references c.f., Sha et al., 2016).

### 4.5. Manipulanda (toy) strategies

Another common strategy for providing EE is to give animals manipulatable “toys.” As defined and categorized here, manipulanda are freely-movable objects or toys that are not part of the cage or feeder and are available to the animals for manual, pedal, oral, or other interaction. These may be toys manufactured for house pets, such as hard rubber cat or dog toys (e.g., Kong™ toys), or those similar to toys manufactured for human children, including dolls, rattles, teething rings, etc. Such manipulanda can vary widely in their complexity and their sensory, motor, and cognitive affordances. In turn, they vary in capacity to elicit different types of engagement. For instance, rattles result in sound production, thus providing motor engagement, sensory stimulation, and feedback loops that may together promote sustained engagement.

There is relatively little evidence that provision of manipulanda or toys have substantial benefits for animal wellbeing. A handful of studies evaluated the effects of manipulanda on abnormal behavior, species-typical behaviors and activity budgets in rhesus macaques (Line, Morgan, & Markowitz, 1991, pigtail macaques (Kessel & Brent, 1998), baboons (Brent & Belik, 1997; Heinz et al., 2000), chimpanzees (Brent & Stone, 1998), and vervets (Alexander & Hines, 2002). In aged rhesus macaques, Line and his colleagues report that simple toys do not alter behavior (Line et al., 1991). In contrast, Brent and Belik (1997) report that provision of manipulanda to group-housed baboons resulted in a decrease in abnormal behavior, inactivity, cage-directed, and self-directed behaviors. A similar result was reported in individually-housed pigtail macaques, with decreases in abnormal behavior and cage-directed behavior correlating with increased enrichment use (Kessel & Brent, 1998). The studies also found that engagement with manipulanda may depend on a multitude of variables, such as, individual preferences, sex preferences, complexity of object, number of toys presented simultaneously, and time of day (Alexander & Hines, 2002; Brent & Stone, 1998; Heinz et al., 2000). Together, the results of these studies demonstrate variability in the use and benefit of using manipulanda as an enrichment strategy to increases animals’ activity and decrease the display of abnormal behaviors.

### 4.6. Mirror strategies

The use of mirrors as EE occurs through provision of either fixed or manipulable reflective surfaces that are not a part of the structure of the animal’s home enclosure, or that are not intended for another purpose. The efficacy of mirrors as an EE strategy has been investigated in vervets (Harris & Edwards, 2004), capuchins (Marchal & Anderson, 1994), and rhesus macaques (Gallup & Suarez, 1991; Suarez & Gallup, 1986). These studies recorded contact use of the mirror, manipulation of the mirror, and duration of viewing behavior exhibited towards the mirror. For example, in singly-housed vervets, mirrors were used up to 26.7% of the time during the observation periods (Harris & Edwards, 2004). Mirror use was defined as either looking behavior directed at the mirror or direct contact with the mirror. Individual monkeys varied in their use of the mirror, but the researchers concluded that the mirrors were a positive and time-consuming enrichment strategy with minimal habituation observed up to a year after introduction (Harris & Edwards, 2004). In contrast, others have reported a cessation of social behaviors directed toward mirrors with extended use (Gallup & Suarez, 1991; Suarez & Gallup, 1986). However, reinstatement of engagement has been shown to occur by altering the location of the mirror (i.e., moving the mirror to a different side of the cage; Suarez & Gallup, 1986) or when the mirrors’ orientation was changed (Gallup & Suarez, 1991). In a study involving eight capuchin monkeys and a variety of mirror types and sizes, the mirrors were manipulated substantially more than the control object (Marchal & Anderson, 1994).

These studies illustrate that when mirrors are available for NHP, they elicit manipulation and engagement. The evidence of manipulative engagement suggests the monkeys could be using the mirrors as manipulanda or as tools to view stimuli that are outside of their direct line of sight (Harris & Edwards, 2004). However, these studies focused on usage and engagement and did not identify other behavioral effects, such as impacts on activity budgets and abnormal behaviors.

### 4.7. Television strategies

Providing EE via television includes presentation of a video monitor screen (without the option for control by the animals) that plays videos of varying content for the animals to watch. As defined here, the category includes provision of video stimuli regardless of specific electronic apparatus used for its delivery (i.e., includes traditional televisions, portable computing devices, etc.). The benefit of providing televisions as an EE strategy has been investigated through studying the success of videos as positive reinforcement, examining the frequency and duration of monkeys’ looking behavior towards the television, and prevalence of species-typical behaviors while television is present in rhesus macaques (Harris, Briand, Orth, & Galbicka, 1999; Platt & Novak, 1997; Schapiro & Bloomsmith, 1995) and pigtail macaques (Lee, Yi, & Crockett, 2011). Platt and Novak (1997) found that monkeys did look at the monitor but for varying lengths of time depending on the content. Similarly, attention to the video screen was found to be higher initially when monkeys were shown a video versus a blank screen, however, rapid habituation and a decline in looking behavior were also observed (Lee et al., 2011). Lee and colleagues found no clear effects on behavior. In a study that allowed monkeys to press a lever to receive video reward, television was a weak positive reinforcement (Harris et al., 1999). Overall, the results of these studies provide no strong conclusion on the efficacy of television as an EE strategy.

### 4.8. Radio strategies

Auditory enrichment can occur in a variety of ways, but one common strategy is to use some form of sound device (without option for control by the animals) to play varied music, talk, conspecific vocalizations, nature sounds, or other streaming auditory stimuli. Radios, compact discs (CD) players, computers that play conspecific sound clips are all grouped here under “radio.” For the purposes of this paper, we will separately consider provision of radio that results from operation of manipulanda and that is paired with food reward (i.e., see section on strategies that combines multiple forms of enrichment). The effectiveness of radio as an EE strategy has been studied in chimpanzees (Howell, Schwandt, Fritz, Roeder, & Nelson, 2003) and baboons (Brent & Weaver, 1996). These studies investigated behavioral changes and changes in heart rate, blood pressure, and plasma cortisol levels. In chimpanzees, stereo music reduced agitation/aggression and increased social behavior after exposure to it (Howell et al., 2003). In baboons, a significant reduction in the mean heart rate during periods when radio was played was interpreted by the authors as an indicator of reduced stress (Brent & Weaver, 1996). The results from these studies suggest that providing music as a source of enrichment may have positive behavioral and physical effects. However, it is difficult to ascertain whether the actual benefit of music is a result of the music itself or the masking “white noise” effect of the music, as engagement with radio has yet to be adequately examined with direct study (Brent & Weaver, 1996).

### 4.9. Compound enrichment strategies

Compound enrichment strategies are those that involve the simultaneous presentation of two or more forms of different types of enrichment. In reality, compound enrichment strategies are likely the current default and common practice. For instance, animals may have toys and mirrors within their housing space, but also be presented with foraging devices and have radios playing. In this section we review those studies that explicitly measure the effect of compound enrichment strategies.

Several studies evaluate the effects of presenting multiple enrichment strategies (typical foraging devices, manipulanda, food puzzles, mirrors, television, etc.) at the same time. The species included in these studies are chimpanzees (Brent & Stone, 1996), rhesus macaques (Griffis, Martin, Perlman, & Bloomsmith, 2013; Line, Clarke, Markowitz, & Ellman, 1990; Schapiro & Bloomsmith, 1994, 1995; Schapiro, Suarez, Porter, & Bloomsmith, 1996), cynomolgus macaques (Bryant, Rupniak, & Iversen, 1988; Turner & Grantham II, 2002), capuchins (Boinski, Swing, Gross, & Davis, 1999; Westergaard & Fragaszy, 1985), and rhesus and cynomolgus macaques (Cannon, Heistermann, Hankison, Hockings, & McLennan, 2016). The range of outcome variables that have been investigated include: cortisol level changes, use of enrichment items, and behavioral effects including changes in activity levels, self-grooming, social interactions, and stereotypical behaviors.

Visits to the “play pens” or supplementing the NHPs’ home cages with enrichment resulted in increases in species typical behavior such as playing, locomoting, and feeding behavior (Bryant et al., 1988; Schapiro & Bloomsmith, 1994, 1995; Schapiro et al., 1996). Decreases in abnormal behaviors, such as stereotypies, were also observed in enriched environments (Cannon et al., 2016; Griffis et al., 2013). On average, cortisol levels were shown to decrease in NHP provided with enriched conditions (Boinski et al., 1999; Cannon et al., 2016; Line et al., 1990).

In a large study that looked at the effectiveness of television as a strategy in combination with typical foraging devices and manipulanda, the sensory (television) enrichment did not result in increased play behavior compared to the physical and feeding enhancements (Schapiro & Bloomsmith, 1995). However, Schapiro and Bloomsmith did find that overall, enriched animals spent more time playing and less time self-grooming than control animals. Another study that observed television, manipulanda and mirror use simultaneously found that the chimpanzees continued to use all three enrichment items over several years (Brent & Stone, 1996). Brent and Stone concluded that television was the most utilized enrichment item based on frequency of looking behavior but found that the chimpanzees varied in the length of time spent looking at the TV monitor depending on the age of the ape (Brent & Stone, 1996).

In study with a novel combination of EE, an enrichment device included a lever that played music and dispensed a pellet when pressed. Singly-housed rhesus macaques consistently engaged with the device, exhibited decreased self-injurious behavior, and trended toward reduced plasma cortisol levels (Line et al., 1991). The option for control along with the provision of food provides a compound stimulus and makes it difficult to differentiate whether the animals are exerting control to gain access to the treats or music itself. Furthermore, most of enrichment plans use radio without the element of control or associated food reward.

Together these studies provide evidence that enhancing environmental complexity using multiple enrichment strategies can promote sustained engagement and have beneficial effects (e.g., reduced SIB). As with other studies of EE, however, the longevity of these effects has not been well studied. One experiment that did evaluate whether the effects of multiple enrichment strategies were sustained after the enrichment was removed reported that the positive effects (i.e., reductions of grooming, stereotypic movement, and self-manipulation behaviors) did *not* persist following their removal (Boinski et al., 1999). The study did not, however, include examination of reintroduction of the enrichment to evaluate whether the previous positive effects were reinstated. It is also important to note that many of the studies that examined a single enrichment strategy likely had other enrichment devices remaining from the animals’ baseline, normal cage environment. In other words, it is unlikely that the specific EE under study was presented in absence of EE typically provided to the animals as part of the facility’s normal procedures. This caveat, in addition to the variable housing conditions between laboratories, make the drawing of general conclusions difficult.

### 4.10. Summary of focused literature review

The evidence reviewed here illustrates how the existing literature can be organized to provide strong empirical basis from which to assign values (or weights) to common EE strategies used in a broad range of settings. The review is not exhaustive, but rather is focused on the types of enrichment included in our assessment tool as guided by the theoretical framework informed by psychological science and described in our companion paper. The broad hypotheses based in this theoretical framework revolve around the sensory, motor, and cognitive affordances – and, hence, stimulation – provided by different EE strategies. In general, the model makes the straightforward prediction that enrichment with more complex and diverse affordances will result in a higher level of engagement, stimulation, or both, and, in turn, will be more likely to benefit NHP wellbeing.

The focused literature review above provides relatively strong evidence that some forms of enrichment elicit high levels of engagement, have the capacity to sustain such engagement, and are associated with benefit to animal wellbeing as indicated by activity levels, reduced stereotypies, absence of SIB, reduced cortisol levels, and so on (although see caveats discussed above). As such, the review highlights empirical evidence for key elements of the assessment tool. For instance, there is clear support for the assignment of a higher value to computer-based enrichment and lower value to passive enrichment such as provision of radio or television. At the same time, the review provides an illustration of two key points: First, how the systematic framework can be used to identify important gaps in knowledge that should be prioritized for further study; and, second, how the tool can be revised and refined with additional evidence.

#### 4.10.1. Species

Overall, review of the extant literature indicates that research to evaluate the effects of EE on NHP is primarily focused on the most common species that are housed in large numbers in a variety of captive settings: macaques, baboons, and chimpanzees. Studies of other apes (gorillas, orangutans), most New World monkeys, and most prosimian primates are relatively few. The disparity likely also reflects the fact that these species are typically in a limited range of captive settings – primarily zoos – and are limited to few individuals of the species within any particular facility. Although research occurs in zoos, it is not the primary focus and thus, may not be a primary source of empirical studies focused on evaluation of different environmental enrichment strategies.

Species differences may result in differential effectiveness of enrichment strategies, thus gaps in knowledge about the use of enrichment and its effects on behavior or wellbeing by some species could pose a challenge to the “fit” of recommendations based in evidence from a limited number of species. In the model proposed here, however, the assessment tool does not differentiate between species, but rather offers a broad framework with inherent flexibility to permit tailoring based on species-specific needs or evidence. To do this, the weights for different factors, or characteristics of enrichment, can be adjusted to correspond to species-typical sensory, motor, or cognitive capacities. Likewise, characteristics may be removed or added. For instance, tailoring the rubric to best reflect the needs of a species that engages in low levels of manipulation, but high levels of olfactory investigation, could be achieved by giving higher weight to enrichment strategies that promote olfactory stimulation and lesser weight to those that require complex manipulation. The process of tailoring the assessment would also highlight priority areas for hypothesis-driven study to determine which enrichment strategies engage the species and which result in benefit to animal wellbeing.

#### 4.10.2. Limitations

Several common characteristics of studies within the domain of EE assessment can be viewed as limitations. In general, such studies focus on relatively short time windows for evaluation of the effects of enrichment. This is true of the time in which behavioral effects are directly measured and the timespan for evaluating the persistence of any changes in behavior associated with the provision of an enrichment strategy. The amount of time that animals engage with an enrichment device, or the presence/absence of abnormal behavior, is often measured by direct observation by a human observer using a 5-10 min period repeated once or twice daily over 1-4 weeks. In general, the persistence of the effects of enrichment on behavioral or wellbeing outcomes are measured in the short-term, over weeks, not months or years. The effect of providing enrichment, removing it, then reinstating is not well-studied.

Finally, few studies include comprehensive measures of wellbeing. Many studies take a “clinical-treatment” approach, seeking to document the effect of EE in terms of reducing abnormal behaviors that include rocking, pacing, self-directed, and self-injurious behavior. Behavioral measures, rather than assessment of physiological, immune, or neural function, predominate in these studies. Arguably, the strongest evidence would come from studies that incorporate multiple measures, across different systems involved in health and wellbeing. The dominance of behavioral observations, however, likely reflects a variety of practical and economic challenges. Obtaining brain images, sampling stress hormones, measuring immune function markers are all associated with a range of costs. Given that there are few sufficient research funding streams associated with scientific study of NHP wellbeing and evidence-based environment enrichment it is not surprising that the relatively non-invasive, unobtrusive, and lower-cost short-term behavioral measurements predominate in the literature.

The limitations and gaps in knowledge in the current body of literature on EE can pose challenges to building an evidence-based program to support NHP wellbeing and to the development of assessment tools. The approach we have taken here, however, offers a framework that not only highlights priority areas for research, but that also allows for modification and strengthening by incorporating the results of continuing empirical study. As additional evidence accrues with greater duration of sampling, longer periods of assessment, and a fuller range of measures, that evidence can be used to adjust the assessment tool in a manner that supports continuing refinement of EE programs.

## 5. Scoring system for assessment of commonly-used environmental enrichment for nonhuman primates

The quantitative assessment tool (or scoring rubric) shown in Table 2 reflects our integration of a theoretically-informed framework along with empirical evidence to construct a workable standard system for evaluation of the relative value of the SMC component of EE programs. As noted in our introduction, there is currently no common systematic approach to evaluate of the NHP EEP that are mandated by federal law. Our major goal here was to devise a relatively straightforward system for categorizing diverse enrichment strategies within a meaningful framework. Thus, we adopted a fairly simple system to minimize ambiguity in categorization and the assignment of points. In all cases, our framework is driven by the existing empirical literature (reviewed above), along with theoretical principles derived from psychological science. Our companion paper provides comprehensive coverage of the latter.

### 5.1. Categories of EE

Within this system, up to three points are assigned for each of the major categories: cognitive, motor, and sensory EE. Each is scored as: absent (0); some (1); and highest level (3). The 0-3-point system reflects an assessment of the degree to which a strategy affords cognitive, motor, or sensory engagement and stimulation. For example, in terms of cognitive stimulation and active engagement, a videogame cognitive test would be assigned a high value (3), while provision of a passive television exposure would be assigned a low value (0).

### 5.2. Characteristics of EE

We also defined cross-category general characteristics of EE that are used for classification and are also assigned points. These include: duration of engagement, choice vs. required, and response contingency. Two characteristics – choice vs required exposure and active vs passive – are scored as present (1 point) or absent (0 points). Categorization of each of these is straightforward. For example, determining whether the animal chooses to engage can readily be assessed by whether their actions result in contact with the enrichment. For a radio playing in the room, the animal is exposed (i.e., required). For a foraging toy, the animal may choose to interact with it, or not (i.e., choice). Similarly, response contingency can be readily and objectively determined by simple consideration of the physical features of the stimuli. Stimuli that change in response to an animal’s action are coded as response-contingent, regardless of the complexity of the response. That is, toys that move, foraging objects from which food can be extracted, videogames that increase in level of difficulty, are all categorized as response-contingent. By contrast, a television or radio is not categorized as response-contingent because the animal’s behavior does not result in a change in the device or the presentation of stimuli.

The third major characteristic and one that is often an outcome measure in evaluation of EE is duration of engagement. Whether an animal engages with enrichment stimuli or not is often taken as a marker of effectiveness. That is, if an animal fails to interact with enrichment at all – doesn’t pick up a toy or avoids a foraging device – it is unlikely that it is an effective or appropriate enrichment strategy. Intuitively, longer duration of engagement would appear to be a positive feature of a strategy if the duration is also associated with positive benefit (i.e., the long duration is not accompanied by negative responses, abnormal behavior, or evidence of aversion). The question of “how long” or what duration is optimal is difficult to address. There are few studies that use continuous measures that would quantify sustained engagement with enrichment devices over periods of hours, days, weeks, or longer. Rather, most of studies calculate percent of time within relatively short observation windows for an individual session (i.e., 5-20 min). Thus, an objective, non-arbitrary categorization of the relative value of duration is not possible based in the current literature. We therefore assigned values based on the existing literature, categorizing duration in terms of 0 (none), 1 (very short, less than 20 min), and 2 (greater than 20 min). The latter category, 20 min or greater, could clearly be further refined with the results of additional empirical study that directly assesses duration of engagement and its effect on behavior and wellbeing. The results of such studies would provide much-needed evidence to determine optimal zones for duration of engagement.

Optimal duration of engagement could occur in a variety of ways. For instance, it could be calibrated to simulate the time budget of animals in field settings where 40% of the animals’ waking time is spent in foraging (Goldstein & Richard, 1989). To mirror this type of engagement, an enrichment strategy would have to elicit sustained attention and manipulation over a 5hr period (see Figure 2A). Alternatively, it may be that relatively brief engagement that occurs consistently and in multiple episodes across a day or over many days may also be optimal with respect to benefit for the animals’ wellbeing (see Figure 2B). Within existing practices, there are enrichment strategies that result in both. For example, our previous studies have demonstrated that monkeys will continue interaction with video games throughout a 5hr period (Bennett et al., 2016) and that engagement with a broad range of different foraging devices typically occurs for briefer periods (20 min; Bennett et al., 2014). In recognition of gaps in the literature, the proposed rubric conservatively categorizes duration of engagement.

**Figure 2.**
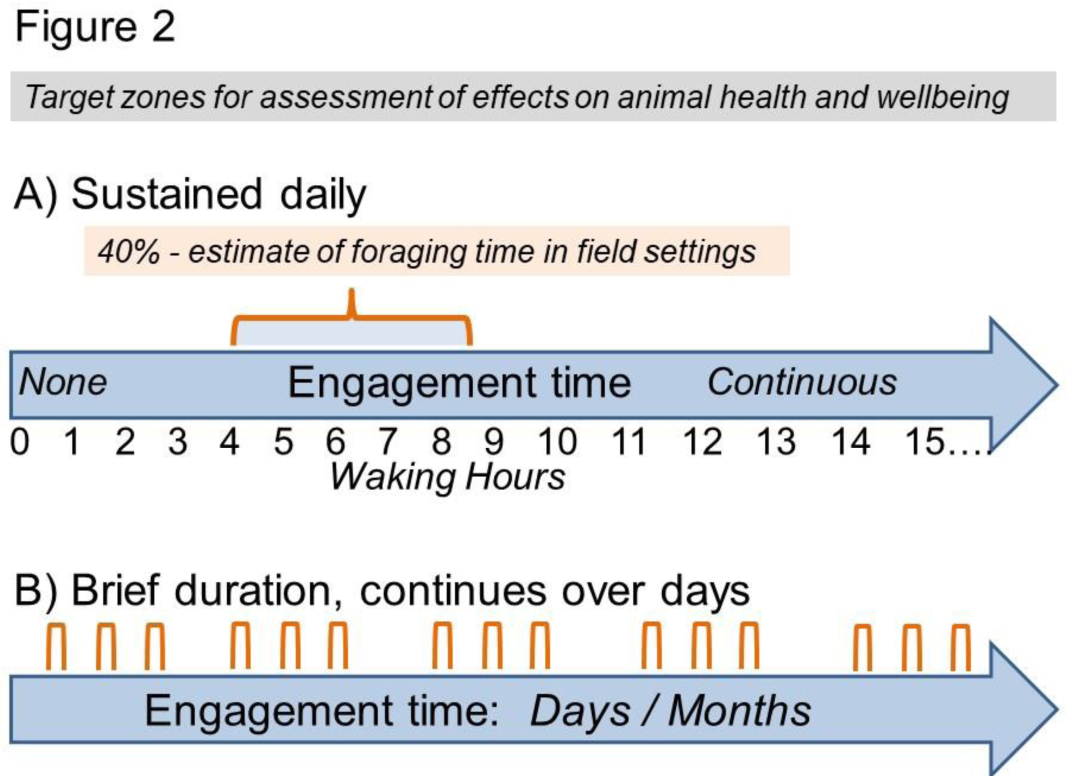
Illustration of time dimensions, patterns of usage of environmental enrichment. A) Sustained engagement over a 24hr period and comparison to daily time budgets for foraging in field settings; B) pattern of brief, but repeated engagement over days or longer periods.

The remaining two characteristics in this rubric are included because they are strong core components of facility managers’ decisions about EE. The presence of a dietary component (i.e., food, treats) and variability, or rotation of devices, each depend heavily on implementation practices within facilities. For instance, captive animals’ total caloric intake is often a clinical or research consideration that influences enrichment decisions. Variation, or rotation, of devices is influenced by several practical and economic factors that include the cost of labor to rotate devices or the cost of maintaining a large and varied inventory.

Anecdotally, both the dietary component and the variability of enrichment receive considerable attention from those charged with devising enrichment plans. Assessing the benefit of each is difficult however, as there is little direct evidence with parametric analysis that would guide categorization. Further, there are no accepted standards for the optimal or expected level of variability in enrichment devices or strategies. Variability can be accomplished by changing the device’s location, contents, or through variations on the device type itself such as incorporating multiple types of puzzle feeders or manipulanda. However, assessing the degree of variability via EEP is difficult and the degree of variability required to observe benefit is not well defined.

Considering these considerations, both dietary component and variability are included in the rubric as present or absent and but do not contribute to the overall score. Their inclusion is simply in acknowledgement of considerations that are important from a practical perspective and that may be important to facility-level decisions that are influenced by benchmarking to community standards. Further, given that specific information about dietary components of EE (i.e., specific foods, variation, frequency of provision) and variability of EE (i.e., how many different types of a device are used, how often they are rotated) is frequently not included in facilities’ EEP, including these components as scored elements within the rubric would likely diminish its utility for cross-facility comparisons. Therefore, each of these considerations points to a gap in knowledge that may be a priority for empirical study.

### 5.3. Overall score

The final calculation step illustrates how this system (derived from the research literature) can be used to examine enrichment strategies and make distinctions among them in terms of their value to animal welfare. We chose the multiplier (or weight) of each category and characteristic according to the strength of empirical evidence and concordance with the theoretical principles from psychological science. Thus, a high weight of 4 is assigned to cognitive engagement because this accords with the empirical evidence of benefit. By contrast, although sensory stimulation is important, as employed in EE it has far less evidence of benefit to animal welfare and therefore is weighted lower with a multiplier of 2. Each of the characteristics (i.e., duration of engagement, choice, response contingency, active/passive), is weighted at 1 (undifferentiated). At present there is insufficient evidence to support a clear differentiation, thus we elected not to assign point values. As with other aspects of the rubric, however, the system allows for modification to accommodate and reflect new evidence, different viewpoints, or priorities.

In Table 4 eight common EE strategies are scored on each dimension, with the total possible points for each category, characteristic, and strategy calculated. As shown, and in accord with the literature reviewed above and elsewhere, the enrichment strategies range from a possible 2-point sum (radio without option for control) to a high value of 29 for both puzzle feeders and joystick or touchscreen cognitive tasks. Furthermore, the proportion of points for each major category reflects its relative value, with the high value (48 points) for cognitive engagement and low value (28) for passive sensory stimulation alone. It is essential to note that cognitive engagement includes elements of sensory stimulation (i.e., visual stimuli presented onscreen, auditory stimuli that accompany responses, manipulative and tactile activity as part of responses, and, possibly to retrieve food rewards).

The scoring scheme proposed here is based on current evidence and is one approach to assigning relative weights for different components of enrichment. The framework is inherently flexible. That is, as discussed above, with additional evidence the weights and points can be changed to reflect changes in knowledge or understanding of the factors that contribute to animal wellbeing. The value of the proposed framework is that it provides a starting point for theoretically-guided, hypothesis-testing approaches to produce evidence that can be used for quantitative approaches to evaluation of SMC EE programs. It is structured to facilitate a Bayesian approach, that is, the current framework specifies a set of priors that can be modified with additional evidence. For instance, as discussed above, experiments designed to test what the optimal duration of engagement with enrichment is in regard to wellbeing outcomes would yield the requisite evidence to inform the framework and allow for finer discrimination between adequate and optimal duration. At the same time, the framework and accompanying review highlight gaps in knowledge that must be addressed to validate the model.

## 6. Illustration of assessment tool use with NHP EEP from zoos, sanctuaries, and research facilities

To provide a demonstration of how the proposed scoring scheme could be applied to assessment of current NHP EEP, we used it to evaluate the content of existing plans shared from two zoos, five sanctuaries, and eight research facilities. We initially contacted over fifty facilities of different types. Fifteen returned their EEP, most with the provision that the facility’s identity remains confidential and not shared in publication. The purpose of the current report is to propose and illustrate the use of the assessment tool; thus, the data reported here are not intended as representative analysis but rather as an illustration of its use for comparative evaluations.

The EEP for all 15 of the facilities were analyzed to produce summary scores (see Table 3). Points in each category were recorded only if the enrichment strategy was plainly stated in the EEP. The level of detail of the plans was widely variable. For many, the EEP provides information at the molar level (i.e., facility provides foraging devices), with facility-specific standard operating procedures (SOPs) referenced for greater detail (i.e., the specific type of devices, the frequency and duration of presentation). It is therefore highly likely that some, perhaps most, facilities may practice additional strategies or provide enrichment with greater frequency than is explicitly stated in their plan. For instance, puzzle feeders and simple foraging devices differ in value, with puzzle feeders that require more intense manipulation and cognitive engagement scoring higher than simple foraging devices. Yet, for many facilities, the type of foraging device may not be contained in the EEP. Thus, the plan score may be an underestimate.

Overall, scoring the plans was relatively straightforward and produced a range that is not unexpected. All facilities met the standard required by the AWA. The range of scores was 25-133, with only one EEP generating the maximum number of points. The scores and rubric provide a relatively straightforward means of comparison and yield a single score. It must be noted, however, that a high score can be achieved using multiple strategies, or the selection of one or two high value strategies. Whether there is a meaningful difference in terms of animal wellbeing benefit between the different approaches that would result in similar high scores cannot be determined from current evidence. It remains for future study directed at testing this hypothesis to evaluate whether high scores achieved through different combinations of strategies result in differences in outcomes for animal wellbeing. This is not a bug in the proposed system, however, but rather illustrates the value of the systematic framework. In short, it produces testable hypotheses and targeted comparisons whose results can be interpreted within a larger, scientifically-valid context. Further, as discussed above, with additional evidence the framework can then be modified to assign relative value in accord with the findings. For example, if use of multiple strategies (i.e., more is better) is associated with greater wellbeing benefit than is use of a single, high value strategy, then additional factors (or weights) could be assigned to the variability component that currently is coded as present/absent but does not contribute to the score.

To provide a basic categorization, scores were divided into quartiles described as minimum, average, exceeds average, and optimal. Table 5 provides data from assessment of an EEP within each of the quartiles. Comparison of these four facilities, which include the range of sanctuaries, zoos, and research facilities, not only illustrates how combinations of different enrichment strategies contribute to an overall score, but also– and perhaps more importantly– provides an accessible visualization. Such visualization offers a direct comparison of EEP that can inform a facility with respect to alignment of its EEP with those of other facilities. Second, the visualization illustrates how a facility can improve its score in a way that reflects meaningful improvement for the animals. Finally, when shared, the visualization provides the basis for continuing progress to identify community standards and best practices.

## 7. Summary: Value of assessment tool to inform continuing refinement of practices and policies about environmental enrichment for nonhuman primates in captive settings

The evidence-based assessment tool developed here provides a straightforward method for evaluating EEP that can serve as a framework to assist individual facilities in evaluation of their EE program and in decision-making about selection of specific EE strategies. Completing the assessment may help a facility more systematically identify gaps in their program or areas for priority consideration and investment. Also, the assessment may help identify low-value practices, those with little evidence for animal welfare benefit, that can be replaced with higher value practices. Of course, decisions about EE involve a range of different stakeholders within facilities, including scientists, facility managers, veterinarians, engineers, animal care staff, compliance staff, and others. Each set of stakeholders may have different concerns and perspectives that inform their decision-making. At minimum, factors that contribute to decisions include: concern for animal wellbeing and selection of best practices to achieve the desired wellbeing outcomes; cost: benefit consideration; practicality and feasibility of the strategy; risk to animals, staff, and facility. All these also must be balanced with weighing the effects of EE on the activity that serves as the central purpose for the facility. As discussed above, for research, the balance may be with scientific objectives, while for exhibitors, the balance may be with visitor perceptions or visibility of the animals.

Our proposed assessment tool provides a framework that can serve as a basis for discussion amongst diverse stakeholders and decision-makers. The assessment tool allows for flexibility and balance at the level of individual facilities. For example, a facility with resource constraints, which is often the case for sanctuaries, community zoos, and small research facilities, may elect to prioritize lower-cost strategies that enhance cognitive engagement rather than investing heavily in toys or sensory enrichment that has little evidence of effectiveness or engages the animals for only short periods of time. The tool can also guide aspirational planning by setting target ranges for improvement, or refinement of EE over time.

### 7.1. Refinement

More broadly, the assessment tool can advance evidence-based refinement of EE at a community level. Our proposed assessment tool provides a flexible, theoretically-grounded and evidence-based system for categorization of EE and evaluation of the value of different strategies. As a result, it is readily adaptable to different types of facilities that balance EE selection with a range of considerations and constraints that vary with the facility’s purpose, economic, geographic, and other factors. At the same time, the tool allows for calculation of an overall score, or single metric. By doing so the assessment framework permits comparison across facilities and the starting point for developing (or better articulating) community standards. Finally, our assessment tool is flexible and readily adapted to account for new evidence. For example, the “multiplier” used here for a category or strategy should be viewed as a proposed value based in current evidence, one that can serve as a starting point for discussion. Additional evidence also may require new characteristics for the categories.

### 7.2. Limitations and future needs

Our organization of the empirical literature to create our assessment tool also identifies areas that need additional hypothesis-driven research to inform best practices in SMC EE. A relatively restricted number of primate species contributed to the generation of the value for each of the categories and their characteristics. For example, many prosimian species are missing from the empirical literature. Moreover, our analysis is of a limited number of plans. The unevenness in non-social, non-structural (i.e., SMC) EE practices is apparent. Of course, a high level of variation is expected given the variability in the comparison of federal regulation and guidance from major accreditation organizations for research facilities, zoos, and sanctuaries and the survey data on common practices. Variability itself is not a major concern because specific practices are driven by a range of factors that differ across facilities. What is of concern, and the major goal with respect to animal welfare, is that the cognitive, motor, and sensory aspects of EE be evaluated by an evidence-based assessment tool to permit the systematic advancement of EEP that best serve animal wellbeing.

We reiterate that our assessment tool is offered as a starting point in developing a system that facilitates the evaluation and establishment of empirically supported EEP. The absence of community standards and a framework for evidence-based decisions serves as a continuing obstacle to progress in advancing best practices in animal care. Furthermore, it appears that there are few mechanisms by which to avoid substituting ineffective practices for those that have meaningful animal welfare benefit. Since there currently is a lack of specificity in standards for the EEP, it is entirely possible that weak practices such as “sensory” enrichment, provision of low-value manipulanda, and quickly consumed supplemental foods could appear to meet the requirement for an EEP. Although implementing engineering standards for EEP would effectively address this concern, moving away from performance standards would likely also have negative impacts for several reasons. As discussed extensively above, engineering standards do not allow for the flexibility that is required for balancing animal welfare, facility purposes, and the range of factors that vary at the facility level. Furthermore, large gaps in knowledge about the effect of many forms of SMC EE serve as a continuing challenge to the creation of policy decisions that are grounded in evidence.

The assessment tool we proposed provides an organizational framework that can guide identification of areas that need additional hypothesis-driven research. Limitations in this assessment tool reflect, in part, limitations in the current evidence. There is currently a lack of evidence about 1) the sustained effectiveness of specific forms of EE; 2) the value of some forms of EE on the wellbeing of animals without clinical concerns such as abnormal and self-injurious behavior; 3) the effects of EE on neural development; 4) direct comparisons of the relative value, in terms of animal wellbeing benefit, of different enrichment strategies; and 5) comparison of the effect of enhanced cognitive enrichment in animals housed in different types of settings (e.g., large group socially-housed in outdoor settings vs smaller social groups, or pairs, housed indoors). The last point highlights the intersection between the assessment proposed here, which is specific to cognitive, motor, and sensory enrichment outside the 3Ss domain and consideration of those other domains. In other words, a full assessment of EEP requires integration of all the domains. To do so will also require evidence about the interactions between the different domains and how that interaction influences psychological wellbeing.

In summary, the assessment tool developed and described here provides a method for quantifying one domain of the EE program. In turn, it can serve as the basis for consideration of refinement, changes in selection of SMC EE, and development of community standards—or best practices—that transcend facility type. This initial effort to create an organizational framework that integrates theoretical perspectives based in psychological science, empirical evidence on EE, and factors that shape decision-making at a facility level, could provide a strong foundation for continuing and focused evolution of evidence-based policy.

## Acknowledgements

We are grateful to Drs. Gabriele Lubach, Sangeeta Panicker, and Taylor Bennett for helpful discussions of evidence, practices, and policies in nonhuman primate care that informed preparation of this paper. We also appreciate the efforts of our undergraduate student research team members who provided assistance: Alina Dain, Trevor Gauthier, Brooke Meidam, Amanda Novak, Erin Schoenbeck, and Aubrey Waldron. This research was partially supported by the University of Wisconsin-Madison Department of Psychology, Graduate School, Wisconsin Alumni Research Foundation, and NIH grant P51 RROD011106.

